# Microbial networks in SPRING - Semi-parametric rank-based correlation and partial correlation estimation for quantitative microbiome data

**DOI:** 10.1101/526871

**Authors:** Grace Yoon, Irina Gaynanova, Christian L. Müller

## Abstract

High-throughput microbial sequencing techniques, such as targeted amplicon-based and metagenomic profiling, provide low-cost genomic survey data of microbial communities in their natural environment, ranging from marine ecosystems to host-associated habitats. While standard microbiome profiling data can provide sparse *relative* abundances of operational taxonomic units or genes, recent advances in experimental protocols give a more quantitative picture of microbial communities by pairing sequencing-based techniques with orthogonal measurements of microbial cell counts from the same sample. These tandem measurements provide absolute microbial count data albeit with a large excess of zeros due to limited sequencing depth. In this contribution we consider the fundamental statistical problem of estimating correlations and partial correlations from such quantitative microbiome data. To this end, we propose a semi-parametric rank-based approach to correlation estimation that can naturally deal with the excess zeros in the data. Combining this estimator with sparse graphical modeling techniques leads to the **S**emi-**P**arametric **R**ank-based approach for **IN**ference in **G**raphical model (SPRING). SPRING enables inference of statistical microbial association networks from quantitative microbiome data which can serve as high-level statistical summary of the underlying microbial ecosystem and can provide testable hypotheses for functional species-species interactions. Due to the absence of verified microbial associations we also introduce a novel quantitative microbiome data generation mechanism which mimics empirical marginal distributions of measured count data while simultaneously allowing user-specified dependencies among the variables. SPRING shows superior network recovery performance on a wide range of realistic benchmark problems with varying network topologies and is robust to misspecifications of the total cell count estimate. To highlight SPRING’s broad applicability we infer taxon-taxon associations from the American Gut Project data and genus-genus associations from a recent quantitative gut microbiome dataset. We believe that, as quantitative microbiome profiling data will become increasingly available, the semi-parametric estimators for correlation and partial correlation estimation introduced here provide an important tool for reliable statistical analysis of quantitative microbiome data.

## 1 Introduction

High-throughput sequencing techniques, including targeted amplicon-based sequencing (TAS) and metagenomic profiling, provide large-scale genomic survey data of microbial communities in their natural habitats. Collaborative efforts such as the Human Microbiome Project (HMP) (Huttenhower et al., 2012), the Earth Microbiome Project (EMP) (Bahram et al., 2018), the TARA Ocean project (Sunagawa et al., 2015), and the American Gut Project (AGP) (McDonald et al., 2018) give an increasingly detailed picture of relative abundances of operational taxonomic units, their phylogenetic relationships, and gene abundances across diverse ecosystems, ranging from marine, soil, and fresh-water to human-associated habitats albeit at different scales and resolutions. Following the seminal work in Woese and Fox (1977), TAS protocols extract and amplify specific regions in marker genes, such as the 16S rRNA gene for bacteria and archaea, the 18S rRNA gene for eukaryotes, and Internal Transcribed Spacer (ITS) regions for fungi, via universal primers followed by next-generation sequencing. These profiling efforts, together with elaborate bioinformatics processing and normalization work flows (Callahan et al., 2016; Caporaso et al., 2010; Edgar, 2013; Lagk-ouvardos et al., 2017; Schloss et al., 2009) allow low-cost determination of highly sparse relative counts of hundreds to thousands of operational taxonomic units (OTUs) or amplicon sequence variants (ASVs) (Callahan et al., 2017; Edgar, 2016) per sample across a large number of sample sites or participants. Metagenomic profiling (Handelsman, 2004) on the other hand provide unbiased samples of the majority of genes of the sampled habitat by high-throughput shotgun sequencing. Sophisticated reference-guided as well as reference free metagenomic read assembly, binning, and taxonomic profiling pipelines (Alneberg et al., 2014; Sczyrba et al., 2017; Sedlar et al., 2017) can, under suitable conditions on read coverage, disentangle the complex mixture of sequencing reads into entire genomes of the underlying microbes and estimate, as a high-level by-product, relative microbial abundances.

Microbiome community-level analysis tasks, such as quantifying community composition shifts across conditions or associating high-dimensional species compositions and their taxonomic profiles to each other and to environmental or host-associated covariates, require statistical estimation procedures that can handle the restrictive nature of such sparse proportional (or compositional) microbiome datasets (Li, 2015). Important examples include differential abundance techniques (Mandal et al., 2015; McMurdie and Holmes, 2014), proportionality estimation (Quinn et al., 2017), regression models with compositional covariates (Holmes et al., 2012; Lin et al., 2014), composition-adjusted correlation estimation techniques (Cao et al., 2018; Friedman and Alm, 2012), and sparse graphical models for microbial association networks (Kurtz et al., 2015; Tipton et al., 2018).

Recent advancements in microbiome profiling protocols, however, promise to alleviate the experimental shortcomings of standard TAS or metagenomic experiments by enabling a more quantitative picture of microbial communities. The experimental protocols in Gifford et al. (2011); Satinsky et al. (2013), originally introduced for marine microbiome profiling, establish quantitative count measurements of environmental metatranscriptomic or metagenomic data by adding orthogonal internal genomic mRNA or DNA standards (of known quantity) to the environmental sample prior to sequencing. A similar spike-in approach has been proposed for gut microbiome studies in Stämmler et al. (2016). Recent quantitative approaches combine TAS techniques with robust measurements of microbial cell counts, in particular flow cytometry (Props et al., 2017; Vandeputte et al., 2017). These tandem measurements provide absolute microbial count data albeit with a large number of zero measurements due to limited sequencing depth (see Fig. 1 for an overview). Thus far, however, statistical analysis methods for these novel quantitative microbiome data remain largely elusive.

**Figure 1:**
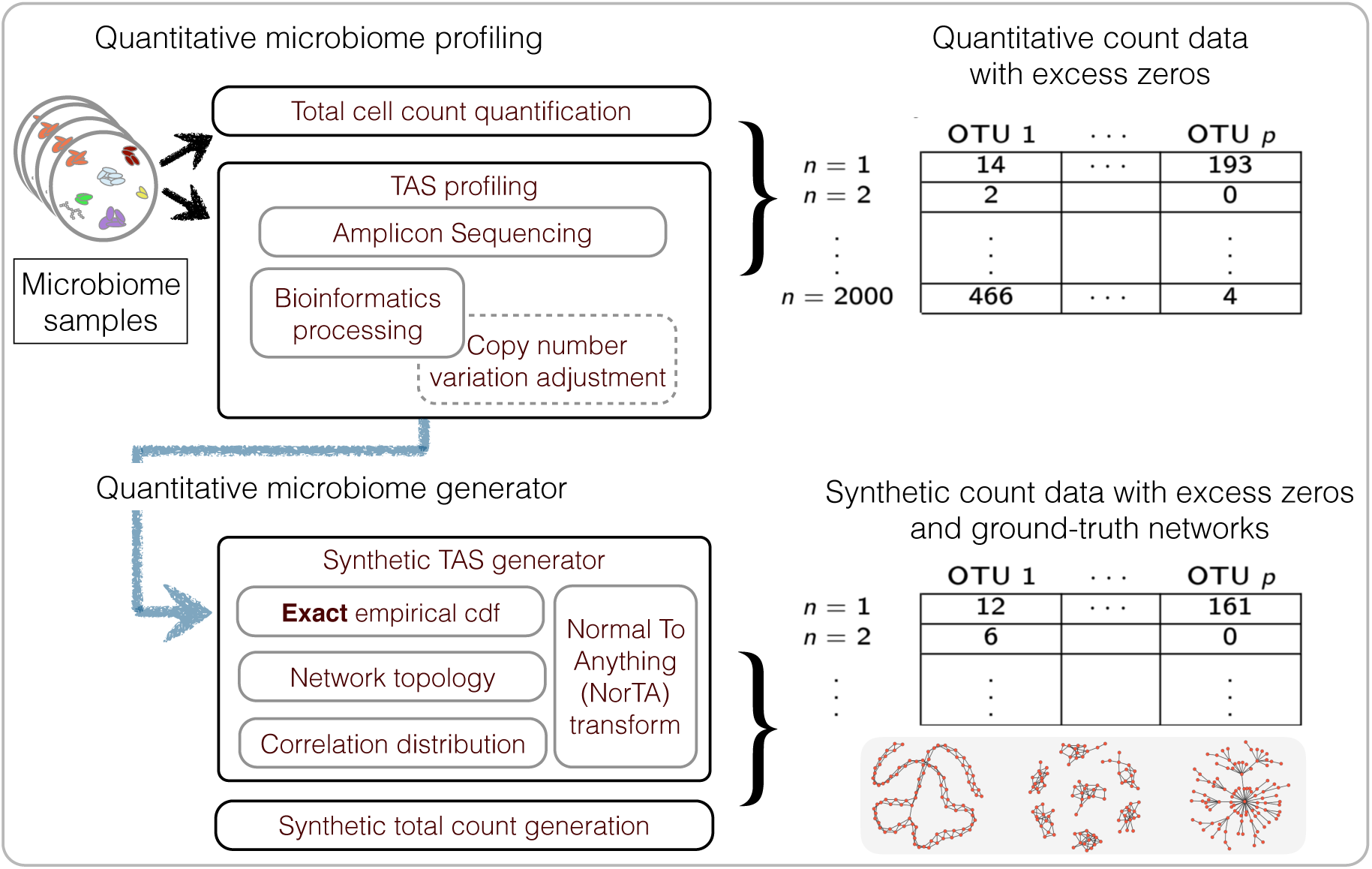
Summary of the workflow for quantitative microbiome profiling (QMP) and the associated synthetic data generation mechanism.

In this contribution, we consider the statistical problem of correlation and partial correlation estimation for sparse *quantitative* microbiome count data. To this end, we first revisit a novel semi-parametric rank-based approach to correlation estimation that can naturally deal with the large number of zeros in the data. This estimator is easy to compute and can readily replace the naïve Pearson or rank-based sample correlation estimator which are often used as a first step in downstream statistical analysis tasks, including principal component analysis, principle coordinate analysis, discriminant analysis, or canonical correlation analysis (Yoon et al., 2018). Here we use the semi-parametric rank-based estimator as a starting point for sparse partial correlation estimation and introduce the **S**emi-**P**arametric **R**ank-based approach for **IN**ference in **G**raphical model (SPRING). SPRING follows the neighborhood selection methodology outlined in Meinshausen and Bühlmann (2006) to infer the conditional dependency graph and uses stability-based model selection (Liu et al., 2010; Müller et al., 2016) to identify a sparse set of stable partial correlation estimates from quantitative microbiome data (Section 2). These partial correlations can be interpreted as direct (i.e., conditionally independent) statistical microbe-microbe associations and can serve as an initial community-level description of the underlying microbial ecosystem (Fuhrman et al., 2015; Ruiz et al., 2017; Sunagawa et al., 2015).

To evaluate our new methodology, we introduce a data generation mechanism that produces synthetic amplicon samples which exactly follow the empirical marginal cumulative distributions of measured amplicon count data while simultaneously obeying user-specified partial correlation dependencies among the variables and closely following user-defined total cell counts (see Fig. 1 for a summary). As ground-truth data for microbial associations remain largely elusive in current literature, our data generation mechanism might be of independent interest for testing other statistical inference schemes. We highlight SPRING’s superior performance compared to standard sparse partial correlation estimation methods on a wide range of quantitative microbiome benchmark problems with varying prescribed network topologies. We also quantify, in the context of association network inference, the potential gains of quantitative over purely relative data even under misspecified totals. To showcase SPRING’s broad applicability (see Section 4), we first infer taxon-taxon associations from relative abundance data collected in the AGP using a pseudo-count-free log-ratio transform that can handle zero counts. Our key application is a genus-level analysis of the quantitative gut microbiome dataset put forward in Vandeputte et al. (2017). We discuss the inferred quantitative association network structure, compare it to published results, and assess, for the first time, the differences between inferred associations from measured absolute and relative abundance data in a consistent statistical framework. While we focus here on TAS-related applications, our methodology is broadly applicable to other data types with excess zeros, including quantitative metagenomics, single-cell RNA-seq, and mass spectrometry data, and thus provides a promising route toward a coherent statistical framework for correlation and partial correlation analysis of multi-omics biological data.

## 2 Semi-parametric rank-based correlation and partial correlation estimation

### 2.1 Rank-based estimation of correlation matrix for zero-inflated data

A great number of multivariate statistical methods, such as principal component analysis, discriminant analysis, canonical correlation analysis and graphical lasso to name a few, require the estimate of covariance or correlation matrix of variables as one of the inputs. The overwhelming number of methods are based on the Pearson sample covariance matrix, which works well at capturing dependencies between variables that are normally distributed. One of the key challenges in analyzing TAS-based microbial abundance data is that it is far from normal: TAS-based measurements are inherently proportional, extremely right skewed, overdispersed, and comprise a large number of zero values. Furthermore, the zeros are not always indicative of the absence of the species, but rather a result of limited sequencing depth or primer bias. For these reasons, the sample covariance matrix is not appropriate for capturing dependencies present in microbiome data. Several methods use techniques from compositional data analysis (Aitchison, 1983), including log-ratio transforms, to adjust the data prior to any estimation, and enforce different structural constraints on the correlation or inverse correlation matrix (Cao et al., 2018; Friedman and Alm, 2012; Kurtz et al., 2015). The problem of excess zeros is typically dealt with by adding an arbitrary pseudocount or, more recently, estimating the pseudocounts from multiple samples (Cao et al., 2017). For quantitative microbiome data, however, correlation and inverse correlation estimators are not yet available. In this work we propose to take a different approach relying on the recently proposed truncated Gaussian copula framework (Yoon et al., 2018).

First, we review the Gaussian copula model, which is sometimes referred to as nonpara-normal (NPN) model (Liu et al., 2009).

#### Definition 1

*A random vector* **x** = (*x*_1_*, …, x_p_*)^*T*^ *satisfies the Gaussian copula model if there exists a set of monotonically increasing transformations* 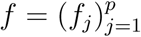 *satisfying* 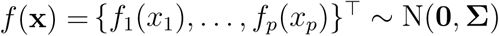 *with σ_jj_* = 1*. We denote* **x** *∼* NPN(**0**, **Σ**, *f*).

The Gaussian copula model is commonly used in undirected graphical models (Fan et al., 2017; Liu et al., 2012) because it models the dependency between variables through the correlation matrix **Σ**, and thus enjoys the mathematical simplicity of Gaussian multivariate distribution while relaxing the normality assumption. While the original model is only appropriate for modeling continuous variables, it has also been generalized to binary variables by adding an extra dichotomization step (Fan et al., 2017). The estimation of graphical models only requires the knowledge of the correlation matrix **Σ**, and it has been shown (Fan et al., 2017) that consistent estimates of **Σ** could be easily obtained from sample Kendall’s *τ* without the need to estimate unknown transformations *f_j_*.

The Gaussian copula model is, however, not appropriate for quantitative microbiome data as (i) it does not take into account zero inflation, and (ii) it models continuous rather than count variables. To address (i), we take advantage of the model proposed in Yoon et al. (2018).

#### Definition 2

(Truncated Gaussian copula model of Yoon et al. (2018)). *A random vector* **x** = (*x*_1_, …, *x_p_*)^*T*^ satisfies the truncated Gaussian copula model if there exists a p-dimensional random vector **u** = (*u*_1_, …, *u_p_*)^*T*^ ∼ NPN(**0**, **Σ***, f*) *such that*

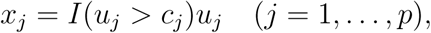

*where I*(*⋅*) *is the indicator function and* **c** = (*c*_1_*, …, c_p_*) *is a vector of positive constants*.

In other words, the model truncates a Gaussian copula variable so it is either zero or positive continuous. This model does not take into account that quantitative microbiome data have zeros or positive counts, but we found the continuous approximation to positive counts to work well in our simulation results (Section 3).

To construct graphical models for the truncated Gaussian copula model, the estimation of the latent correlation matrix **Σ** is required. Yoon et al. (2018) develop a rank-based estimator for **Σ** by deriving the explicit form of the so-called bridge function *F* that connects the sample Kendall’s *τ* estimates to the elements of **Σ**. Given observed data (*x_j_*_1_*, x_k_*_1_), …, (*x_jn_, x_kn_*) for variables *j* and *k*, the sample Kendall’s *τ* estimate is defined as

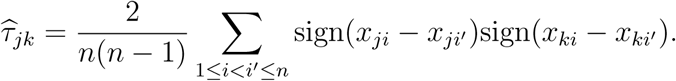

The bridge function *F* is defined so that 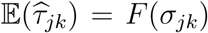, where *σ_jk_* is the corresponding latent correlation between variables *j* and *k*. The explicit form of *F* for the truncated Gaussian copula model is given below.

#### Theorem 1

(Yoon et al. (2018)). *Let random variables x_j_, x_k_ follow truncated Gaussian copula with corresponding latent correlation σ_jk_. Then* 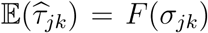, where

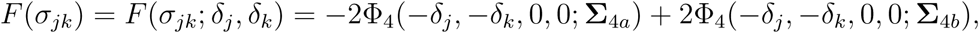

δ_j_ = *f_j_*(*c_j_*)*, δ_k_* = *f_k_*(*c_k_*), Φ_4_(…; **Σ**_4_) *is the cumulative distribution function (cdf) of the four dimensional standard normal distribution with correlation matrix* **Σ**_4_,

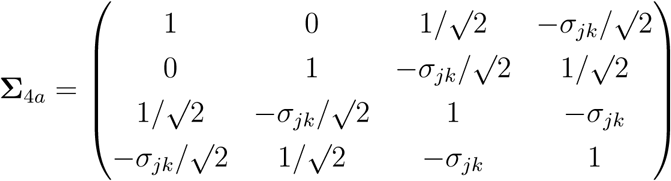

and

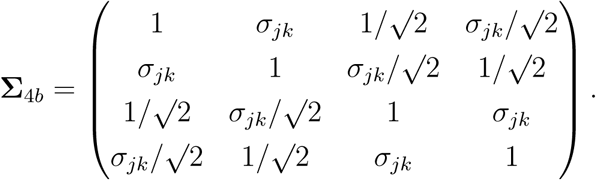

Moreover, *F*(*σ_jk_*) *is strictly increasing, so the inverse function F* ^−1^(*σ_jk_*) *exists*.

Theorem 1 provides a closed form of the bridge function *F* that can be used to estimate *σ_jk_* by calculating 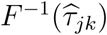. The thresholds *δ_j_* can be easily estimated from the moment equations. In practice, the inverse of the bridge function 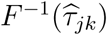 is calculated numerically by finding the minimizer of the quadratic function 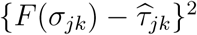, which is unique due to the strict monotonicity of the function *F* (*σ_jk_*). We refer to Yoon et al. (2018) for further estimation details. The entire procedure is implemented within the R package `mixedCCA` (Yoon and Gaynanova, 2018).

In summary, the approach allows the construction of a rank-based estimator 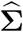 for the correlation matrix **Σ** of truncated Gaussian copula variables, which forms the basis for the undirected graphical model framework in Section 2.2.

### 2.2 Sparse graphical models and SPRING

We next introduce the **S**emi-**P**arametric **R**ank-based approach for **IN**ference in **G**raphical model (SPRING). SPRING relies on the estimation of an undirected graphical model from data. Undirected graphical models are typically used to represent the conditional independence relationship between the variables of random vector **x** *∈* ℝ^*p*^, so that

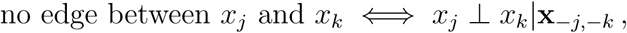

where **x**_*−j,−k*_ means all components in **x** except component *j* and *k*. If the vector **x** follows a normal distribution, then conditional independence between *x_j_* and *x_k_* is equivalent to zero partial correlation between variables *j* and *k*. Therefore, sparse estimates of partial correlations lead to sparse conditional independence graphs. There is a rich literature on sparse estimation of partial correlations, with perhaps the most popular methods being the neighborhood selection of Meinshausen and Bühlmann (2006) (denoted by MB from here on) and the graphical lasso (Friedman et al., 2008). While the rank-based estimator of the correlation matrix proposed in Section 2.1 can be used in both approaches, we found the MB method to perform better than graphical lasso in numerical simulations, and therefore focus on the MB method in the remainder of the paper.

The MB method takes advantage of the connection between partial correlations and regression coefficients and performs sparse estimation of partial correlations by regressing each of the *p* variables on the rest, thus finding each nodes’ immediate neighbors by solving a lasso problem (Tibshirani, 1996). Given column-centered and scaled data matrix **X** *∈* ℝ^*n×p*^ with columns **x**^*j*^, the MB method solves for each variable *j*

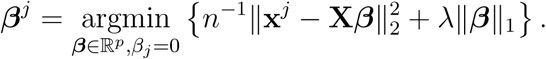

Rewriting the objective function leads to

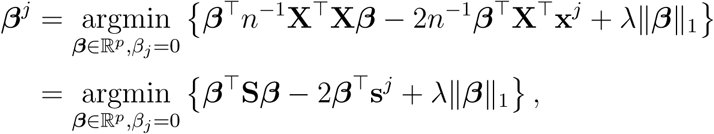

where, given the centering and scaling of **X**, **S** = *n*^−1^**X**^*T*^**X** is the sample correlation matrix with columns **s**^*j*^. Since the standard sample correlation matrix is not suited for capturing dependencies in sparse quantitative microbiome data, SPRING replaces the sample correlation **S** in the MB method with the proposed rank-based estimator 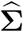 from Section 2.1. The MB method comprises the regularization parameter *λ* which balances the trade-off between sparsity of the neighborhood and goodness of fit, and thus requires data-driven tuning. We here consider a stability-based model selection method, the Stability Approach to Regularization Selection (StARS) (Liu et al., 2010), which has been previously proven to be suitable for graphical model selection on microbiome data (Kurtz et al., 2015; Müller et al., 2016). The StARS method selects the optimal tuning parameter by repeatedly taking subsamples of the original data, estimating the graphical model for each subsample at each *λ* value along a prescribed regularization path, and then calculating empirical edge selection probabilities from the subsamples. The StARS edge stability criterion uses these probabilities to assess the sum of edge variabilities for each graph along the regularization path. The optimal *λ* is selected based on the supplied threshold t_S_, with standard values being t_S_ = 0.05 and t_S_ = 0.1 (Kurtz et al., 2015; Liu et al., 2010). The threshold value represents a bound on the allowed overall edge variability over the entire graph. Lower thresholds lead to sparser, more robust graphs. Using the selected *λ* value, the final graphical model is refitted on the full dataset.

In summary, SPRING comprises three major components: (i) a semi-parametric rank-based correlation estimator for zero-inflated count data, (ii) the MB method to infer sparse conditional dependencies from the estimated correlation, and (iii) a stability-based approach (StARS) for sparse and robust neighborhood selection.

### 2.3 Extensions to compositional data

An important prerequisite for SPRING to be applicable to zero-inflated data is that individual count values across samples are comparable. For TAS-based microbial abundance data this condition is not satisfied because the total read count of a sample is not related to the total number of bacteria in the sample (Vandeputte et al., 2017), thus making the counts inherently proportional quantities. While this drawback is alleviated with the novel experimental techniques for quantitative microbiome data, as discussed earlier, a large number of available datasets, including the HMP and the AGP data, are only available as proportional (or compositional) data. To make SPRING amenable to statistical association inference from relative abundance data, we propose a novel data transformation as follows.

One of the key challenges in working with compositional data is the presence of unit-sum constraint. For correlation estimation, a common approach (see, e.g., Aitchison (1983); Cao et al. (2018); Kurtz et al. (2015)) is to first apply the centered log-ratio transform (clr) to the compositional vector of each sample 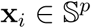

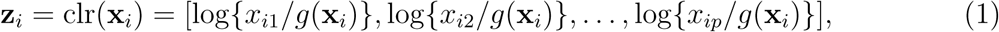

where 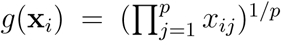 is the geometric mean of **x**_*i*_. A correlation matrix is then estimated based on the transformed **z**_*i*_, *i* = 1*, …, n*, rather than directly on **x**_*i*_ (Aitchison, 1983). Since TAS-based microbiome profiling data have a large number of zeros, the addition of a large number of pseudo-counts is required to modify the vector of compositions to only have non-zero proportions. Adding such pseudo-counts modifies the measured non-zero proportions and masks the zeros in the data, leading to zeros and non-zeros being treated equally in subsequent analysis. In addition, the choice of the actual value of the pseudocount can influence downstream analysis results, and mere addition of extra zero components to the compositional vector would also change the transformation.

To avoid these drawbacks and to play on the strengths of SPRING in handling excess zeros, we propose a modified clr transform (mclr) that avoids the use of a pseudo-count. Contrary to recent efforts in data-driven inference of pseudo-counts (see, e.g., Cao et al. (2017); de la Cruz and Kreft (2018) and references therein), we compute the geometric mean of each sample from positive proportions only, normalize and log-transform all non-zero proportions by using that geometric mean, and apply an identical shift operation to all non-zero components in the dataset. Specifically, let 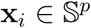 be the vector of compositions for sample *i*, and for simplicity of illustration assume that the first *q* elements of **x**_*i*_ are zero, and the other elements are non-zero. Then we propose to apply

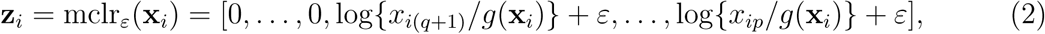

where 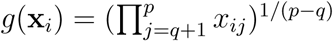 is the geometric mean of the non-zero elements of **x**_*i*_. When *ε* = 0, mclr_0_ corresponds to clr transform applied to non-zero proportions only. When *ε >* 0, mclr_*ε*_ applies additional positive shift. To make all non-zero values strictly positive, we use a data-driven shift *ε* = *|z*_min_*|* + *c*, where *z*_min_ = min_*ij*_ log*{x_ij_/g*(**x**_*i*_)*}* and *c* is any positive constant with default value *c* = 1. The rationale for the shift is to preserve the original ordering of the entries of the compositional vector **x**_*i*_ (with zeroes being the smallest) in the transformed vector **z**_*i*_. The constraint *ε > |z*_min_*|* ensures that *z_i_*_(*q*+1)_*, …, z_ip_* are strictly positive for all *i*. The modified clr transform is invariant to the addition of extra zero components, preserves the original zero measurements, and is overall rank-preserving.

If a practitioner intends to infer microbial associations from relative abundance data using SPRING, we suggest to first use the mclr_*ε*_transform on relative abundance data and then apply SPRING to the transformed data. While SPRING is completely invariant to the choice of *ε* in mclr_*ε*_for any value of *ε* within the constraint, it does not take into the account the compositional nature of the data. Alternative ways of measuring associations between compositional components include Aitchison’s variation (Aitchison, 2003), linear compositional associations (Egozcue et al., 2018), and proportionality (Quinn et al., 2017), which take the compositional constraints directly into account. Here, we will focus on correlation-based approaches and present an application of SPRING to AGP data in Section 4.1.

## 3 Simulation studies

### 3.1 Generation of synthetic quantitative microbial abundance data

Our first goal is to develop a generating mechanism for synthetic data with given conditional independence graph that emulates as close as possible *quantitative* microbial abundance data. We closely follow ideas presented in Kurtz et al. (2015) for synthetic data generation with several important differences. The work flow of our data generation mechanism is summarized in Figure 1.

First, we discuss how to generate the correlation matrix **Σ** given an adjacency matrix **Θ**. Let *p* be the number of nodes, i.e., the number of taxa or OTUs, and let **Θ** be the *p* by *p* adjacency matrix so that *θ_ij_* = 1 if there is an edge between nodes *i* and *j*, *i ≠ j*, and *θ_ij_* = 0 otherwise. We assume the induced graph has no self-loops so that *θ_ii_* = 0. We here consider three types of graph topologies: band graphs, cluster graphs, and scale-free graphs. The number of edges in the graph is denoted by *e*. The default value considered here is equal to twice the number of nodes (*e* = 2*p*), resulting in sparse graphs. Given this fixed sparsity level and the graph type, we use the R package SpiecEasi (Kurtz et al., 2017) to generate a precision matrix **Ω** with the pattern of zeros corresponding to **Θ**. Using **Ω**, we generate the correlation matrix **Σ** by taking the inverse of the precision matrix, followed by scaling.

Given the correlation matrix **Σ**, we follow Kurtz et al. (2015) and use the “Normal to Anything” (NorTA) approach to generate synthetic abundance data. The NorTA method allows to generate variables with arbitrary marginal distributions from multivariate normal variables with given correlation structure. Specifically, we first generate *n × p* matrix **Z** with independent normal rows **z**_*i*_ ∼ N(**0**, **Σ**) with given correlation matrix **Σ**, then get uniform random vectors by applying standard normal cdf transformation to each column of **Z**, **u**^*j*^= Φ(**z**^*j*^) element-wise, and then apply the quantile functions of the target marginal distributions to each **u**^*j*^. In Kurtz et al. (2015), the zero-inflated negative binomial distribution (zinegbin) from VGAM package (Yee, 2010) is used, where the marginal distributional parameters are estimated from measured amplicon data. However, we found that the zinegbin distribution does not emulate well the overdispersion and skewness present in real data. This is evident by comparing the summary statistics between, e.g., the AGP data and corresponding synthetic data generated using the zingebin, as shown in Table 1. To better match real amplicon data, we propose to take a different approach by using the inverse of the *empirical* cumulative distribution function (ecdf) of each OTU. This inverse can be calculated numerically by using the uniroot.all function in rootSolve package in R (Soetaert, 2009). As is evident from Table 1, the ecdf approach works well in mimicking the summary statistics of real TAS-based data. The match across all counts is considerably better than the match across sample abundances since the ecdf transformation is applied separately to each OTU. Although the within-sample counts are affected by the imposed correlation structure **Σ**, the values of the sample total abundance of synthetic data with the ecdf are much closer to the measured ones than those with zinegbin. In terms of count summary statistics, the synthetic data is nearly indistinguishable from the measured data.

**Table 1.** Comparison of summary statistics for all the counts and sample total abundance values between AGP data and two synthetic data generators. The sample size is *n* = 2000, the number of OTUs is *p* = 200, and the synthetic data is based on scale-free graph. type.

| Count data |  |  |  |  |  |  |
| --- | --- | --- | --- | --- | --- | --- |
| Data | Min. | 1st Qu. | Median | Mean | 3rd Qu. | Max. |
| American Gut Project | 0.0 | 0.0 | 3.0 | 144.9 | 34.0 | 176673.0 |
| Synthetic (zinegbin) | 0.0 | 0.0 | 33.0 | 170.0 | 125.0 | 54704.0 |
| Synthetic (ecdf) | 0.0 | 0.0 | 3.0 | 144.3 | 34.0 | 176673.0 |

| Sample total abundance |  |  |  |  |  |  |
| --- | --- | --- | --- | --- | --- | --- |
| Data | Min. | 1st Qu. | Median | Mean | 3rd Qu. | Max. |
| American Gut Project | 10002.0 | 15149.2 | 20715.5 | 28989.0 | 32964.8 | 341632.0 |
| Synthetic (zinegbin) | 21696.0 | 30119.2 | 32543.5 | 33995.7 | 36068.0 | 86033.0 |
| Synthetic (ecdf) | 7354.0 | 18860.5 | 25095.5 | 28854.7 | 34285.2 | 196732.0 |

### 3.2 Synthetic data generation from American Gut Project data

In this paper, we generate synthetic counts from a large subset of the American Gut Project data (McDonald et al., 2018), which comprises *p* = 27116 taxa across *n* = 8440 samples. We first prune these data using the following three steps: (1) exclude samples whose sequencing depths (total read abundances) are *≤* 10, 000; (2) exclude all taxa present in less than 30% of samples; (3) exclude samples whose abundance is less than the first percentile of all sequencing depths. This leads to a reduced dataset with *p* = 481 taxa across *n* = 6482. We consider two scenarios for the simulation studies: a large and a small sample size setting. For the large sample size setting, we randomly pick *n* = 2000 samples with total abundance at least 10,000, and then select *p* = 100 OTUs with largest abundances leading to 2000 *×* 100 matrix of synthetic counts. For the small sample size setting, we use the same strategy with *n* = 500 and *p* = 200. In the synthetic benchmarks, we treat the total observed read abundances as quantitative microbiome profiling abundances and impose the different conditional dependencies on these counts. We refer to these samples as “True data” in the simulations. To investigate the robustness of SPRING to misspecifications of the assumed total, we also generate “Distorted data” by multiplying counts in every sample with an individual scale factor chosen uniformly at random from the interval [0.5, 3]. The scale factor does not affect a sample’s compositional data but does distort the total abundances. The scale factor interval [0.5, 3] represents a realistic distortion scenario in gut microbiome samples (see e.g., in Vandeputte et al. (2017), Figure 2) and is on the same order as typical fold changes of observed image-based total species counts in marine ecosystems (Fuhrman, 2018). We study the performance of SPRING both on the “True” and “Distorted” synthetic data in order to assess how strongly a misspecification of the total affects association network inference.

**Figure 2:**
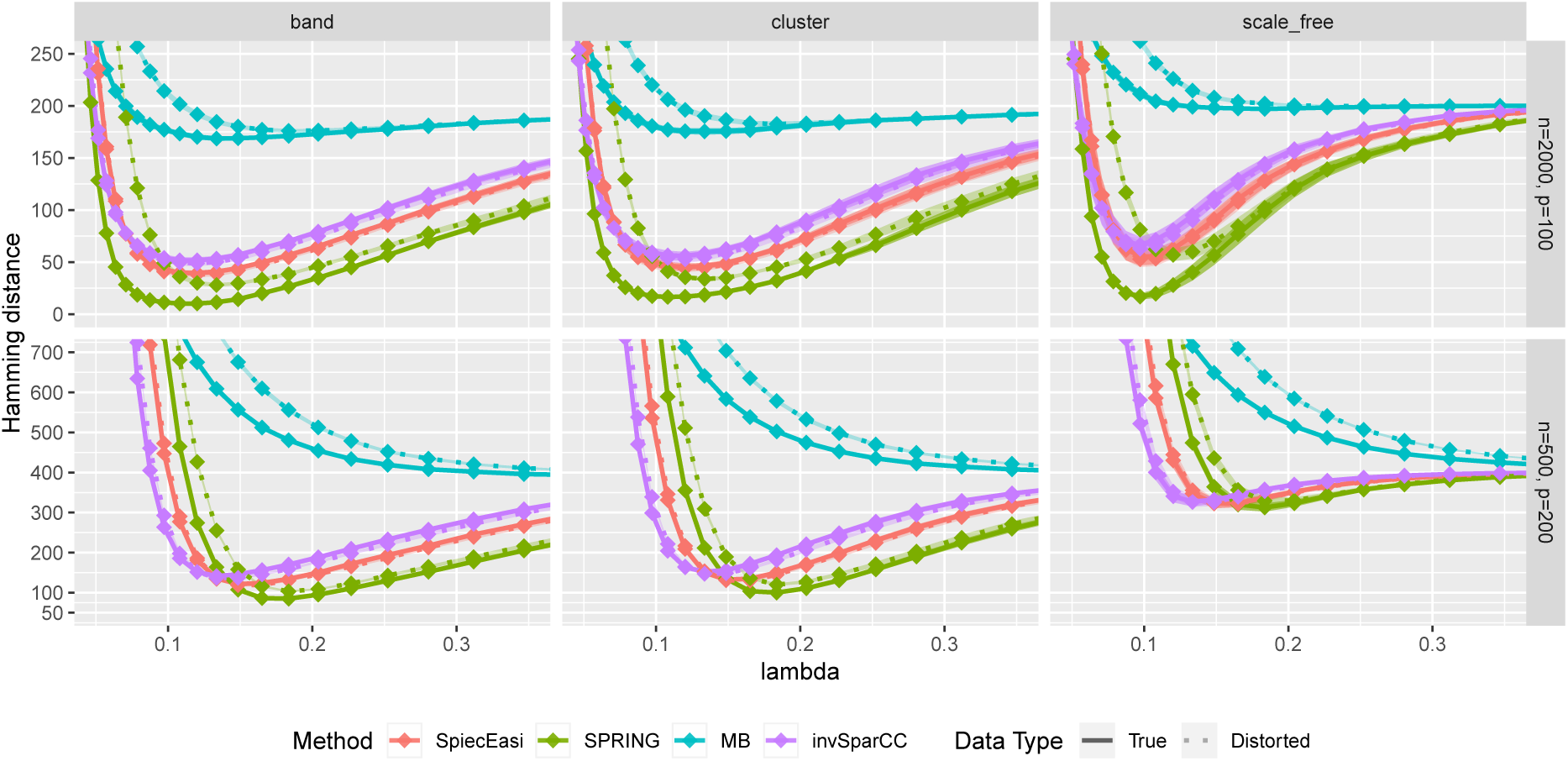
Hamming distance as a function of tuning parameter *λ*. The lines correspond to mean values across 50 replications, and the bands show two standard errors. True abundance and distorted abundance data are distinguished by the transparency level and the line type: true data are less transparent and have solid lines; distorted data are more transparent and have dotted lines.

### 3.3 Methods for comparison

Along with SPRING, we consider three methods of comparison. To study the influence of the sample correlation estimation, we consider the standard MB method using the Pearson sample correlation (Meinshausen and Bühlmann, 2006) (implemented in the R package huge (Zhao et al., 2012)). We also consider two popular methods for microbial association inference from relative abundance data: SPIEC-EASI in the MB mode (Kurtz et al., 2015) and SparCC (Friedman and Alm, 2012) (both implemented in the R package SpiecEasi). The original SparCC method, however, is used for inferring marginal rather than conditional dependencies. For fair comparison with the other methods, we therefore introduce a modification of SparCC, termed invSparCC. The invSparCC method estimates the correlation matrix using the default SparCC method (as implemented in the R package SpiecEasi), and then uses the SparCC correlation estimator as input to the MB method, described in Section 2.2. All considered methods use the neighborhood selection principle to derive a sparse graphical model. The inferred adjacency and coefficient matrices are thus not guaranteed to be symmetric. We use the “or” rule and the “maxabs” rule to symmetrize the estimated adjacency and coefficient matrices, respectively. The “or” rule assigns an edge between nodes *i* and *j* if either node *i* is selected as a neighbor of *j* or node *j* is selected as a neighbor of *i*. The “maxabs” rule symmetrizes the coefficient matrix by taking the coefficient with maximum absolute value. For tuning parameter *λ* selection, we use the R package pulsar with “StARS” edge stability criterion and use 50 subsamples with subsampling ratio being fixed at 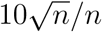, where *n* is the sample size.

### 3.4 Results

We first compare the methods in terms of the Hamming distance between the true and the estimated graph. The Hamming distance is calculated as the number of edges that disagree with the true graph at each value of tuning parameter *λ*. The comparison of Hamming distance curves across the values of *λ* allows us to check the best achievable Hamming distance value that is agnostic to tuning parameter selection scheme. We consider 50 values of *λ* for all methods equally spaced on a logarithmic scale, with *λ*_max_ corresponding to no edges in the estimated graph, and *λ*_min_ = 0.01*λ*_max_. For more accurate comparison, we consider 50 replications of the data generating process for each specified combination of *n* and *p*. The mean Hamming distance values over 50 replications as functions of *λ* are plotted in Figure 2, with bands corresponding to *±* two standard errors. The MB method is uniformly outperformed by all methods, confirming that standard sample correlation is not suitable for capturing dependencies in sparse quantitative microbiome data. SPIEC-EASI and invSparCC have comparable performance, with SPIEC-EASI achieving smaller mean values. SPRING performs best in all cases considered here. The most challenging scenario is the scale-free graph with low sample size, with SPRING, SPIEC-EASI, and invSparCC having comparable performance. As expected, the distortion of total abundances has no effect on the compositional methods SPIEC-EASI and invSparCC, but decreases the performance of MB and SPRING. Nevertheless, the minimum Hamming distance achieved by SPRING on distorted data is still comparable or better than the minimum distances achieved by other methods, thus suggesting that SPRING is robust to misspecification of total abundance values.

Next, we consider one data replication, and compare the Hamming distances achieved by selecting the tuning parameter *λ* using StARS. The results are shown in Figure 3 with two StARS thresholds considered (stars indicating 0.1 and circles indicating 0.05). As expected, smaller threshold corresponds to larger tuning parameter leading to sparser graph. At the same time, based on numerical results, the threshold of 0.1 tends to reach smaller Hamming distances for all methods except MB. In general, both thresholds lead to reasonable values of *λ* in terms of Hamming distance. As in the previous comparison, SPRING leads to smaller Hamming distance values for “True” data and is robust to misspecified total abundance values.

**Figure 3:**
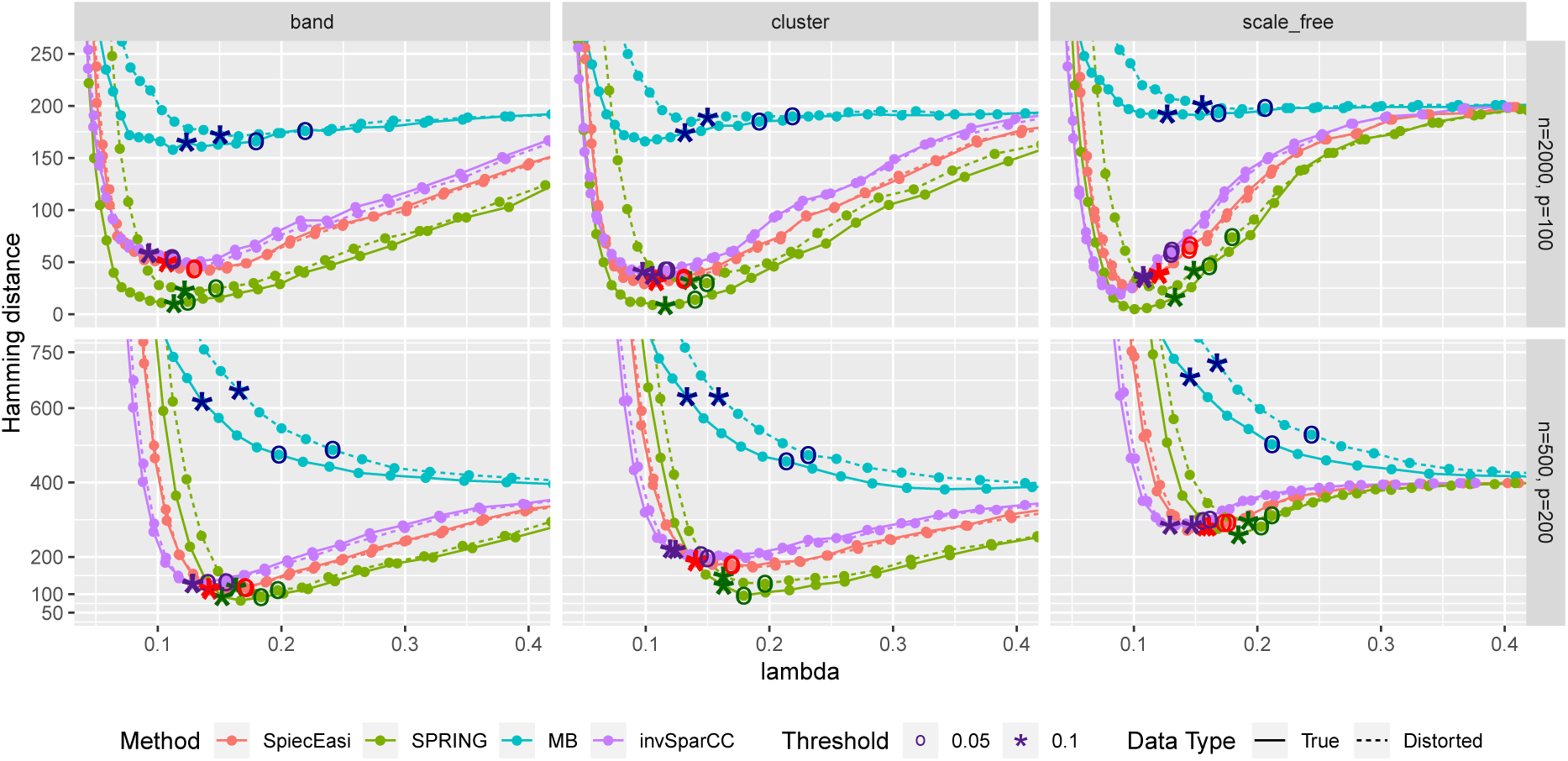
Hamming distance as a function of tuning parameter *λ*. The distances at the tuning parameters selected by StARS are marked with star-shaped points (t_S_ = 0.1) and circle-shaped points (t_S_ = 0.05). True data are plotted with solid lines and distorted data are plotted with dotted lines.

Finally, we compare the estimated graphs from all methods in terms of precision and recall curves, where

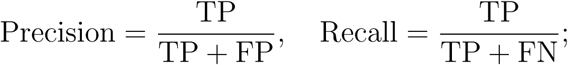

TP, FP, and FN indicate the number of True Positives, False Positives, and False Negatives, respectively. To construct the curves, we extract the edge selection probabilities based on 50 subsamples from pulsar corresponding to tuning parameter with t_S_ = 0.1. We calculate precision and recall values by changing the threshold for edge selection probability from 1 to 0, interpolating the precision-recall values at the edges for no selection (recall = 0, precision = 1) and complete selection (recall = 1, precision = 4*/*(*p −* 1)). Here 4*/*(*p −* 1) is the probability of choosing true edges (*e* = 2*p*) at random among all possible edges (*p*(*p−*1)*/*2). The resulting curves are in Figure 4. For True data, SPRING achieves the highest precision-recall curves across all scenarios. The Area Under the Precision-Recall curve (AUPR) values are reported in Table 2. For the distorted data, SPRING is still best or among the best methods for band and cluster graph types, and is outperformed by the compositional methods for scale-free graph type in the low sample size regime.

**Table 2.** Area under the Precision-Recall curves (AUPR) of Figure 4. In each cell, AUPR of the True data and the Distorted data (given in parenthesis) are reported. AUPR value is based on edge selection probabilities using StARS with t_S_ = 0.1.

| Dimension $(n, p)$ | graph type | SPIEC-EASI | SPRING | MB | invSparCC |
| --- | --- | --- | --- | --- | --- |
| (2000, 100) | band | 0.91 (0.92) | 0.95 (0.94) | 0.42 (0.34) | 0.91 (0.91) |
|  | cluster | 0.93 (0.93) | 0.95 (0.92) | 0.32 (0.27) | 0.93 (0.93) |
|  | scale-free | 0.93 (0.93) | 0.96 (0.93) | 0.26 (0.18) | 0.94 (0.94) |
| (500, 200) | band | 0.89 (0.90) | 0.93 (0.89) | 0.20 (0.16) | 0.87 (0.88) |
|  | cluster | 0.83 (0.84) | 0.90 (0.88) | 0.25 (0.22) | 0.81 (0.82) |
|  | scale-free | 0.55 (0.54) | 0.58 (0.50) | 0.01 (0.01) | 0.54 (0.54) |

**Figure 4:**
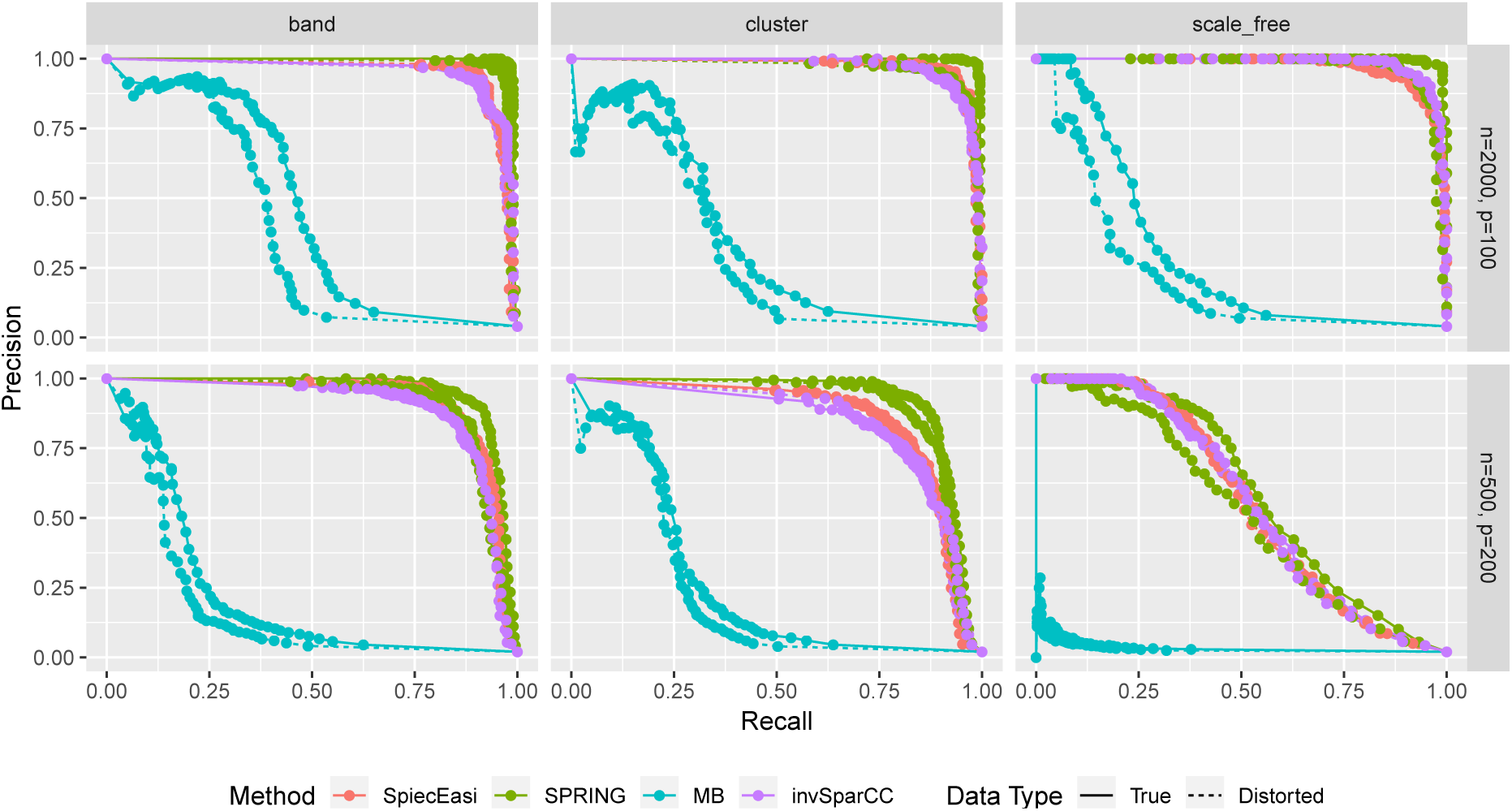
Precision-Recall curves based on edge selection probabilities from pulsar with t_S_ = 0.1. True data are plotted with solid lines and distorted data are plotted with dotted lines.

In conclusion, SPRING exhibits considerably better graph recovery performance than existing methods, and is quite robust to misspecification of total sample abundance. This suggests that incorporating quantitative abundance information in the analysis leads to more reliable graphical model inference.

## 4 Statistical microbial associations in gut microbiome data

We provide two applications of SPRING to TAS-based microbial abundance data: a subset of the relative abundance data from the American Gut Project (AGP) (McDonald et al., 2018) and the QMP data from Vandeputte et al. (2017).

### 4.1 Taxon-taxon associations from the American Gut Project data

We first use SPRING to infer taxon-taxon associations from the relative abundance AGP data. After the pruning and filtering steps described in Section 3.2, we arrive at *p* = 481 OTUs from *n* = 6482 samples. Prior to applying SPRING, we transform the compositions 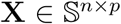 using the mclr_*ε*_ transform introduced in Eq. 2. The minimum value of the mclr_0_-transformed data across all samples is *z*_min_ = *−*4.8142. To make all non-zero values strictly positive, we add an arbitrary constant *c* = 1 to *|z*_min_*|* and use the shift *ε* = *|z*_min_*|* + *c* = 5.8142 in the final mclr_*ε*_ transform. We also consider SPIEC-EASI, MB, and invSparCC (see Table 3) for comparison. All four methods use the same parameterization for the regularization path and StARS model selection: 50 subsamples with the same seed number, subsampling ratio 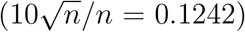 and 50 tuning parameter values with the same ratio of the smallest to largest *λ* value (*λ*_min_*/λ*_max_ = 0.01). For each method, *λ*_max_ is set to the maximum value of the off-diagonal elements of the respective correlation matrix. All computations were performed in R using the R packages `pulsar, SpiecEasi, huge`, and `mixedCCA`, respectively.

**Table 3.** Summary of methods considered for comparison. For all methods, the final graphical model is estimated based on combining neighborhood selection approach with pulsar tuning parameter selection. * - when absolute abundance data is not available, SPRING can be applied to relative abundance data following mclr transform described in Section 2.3.

| Method | Type of data | Transformation | Correlation estimation |
| --- | --- | --- | --- |
| MB | absolute abundance | none | sample correlation |
| SPIEC-EASI | relative abundance | clr | sample correlation |
| invSparCC | relative abundance | log | SparCC |
| SPRING | absolute/relative abundance* | none/mclr* | latent correlation |

We report summary statistics of the estimated association networks for two StARS stability thresholds: 0.05 (the standard setting in SpiecEasi) and 0.1 (the standard setting in Liu et al. (2010)) in Table 4. For both stability thresholds, the MB method estimates the sparsest networks with highest percentage of positive edges (PEP) while invSparCC estimates the densest networks with the lowest percentage of positive edges. SPRING and SPIEC-EASI’s association networks have similar edge densities while SPRING has a considerably higher percentage of positive partial correlation edges.

**Table 4.**
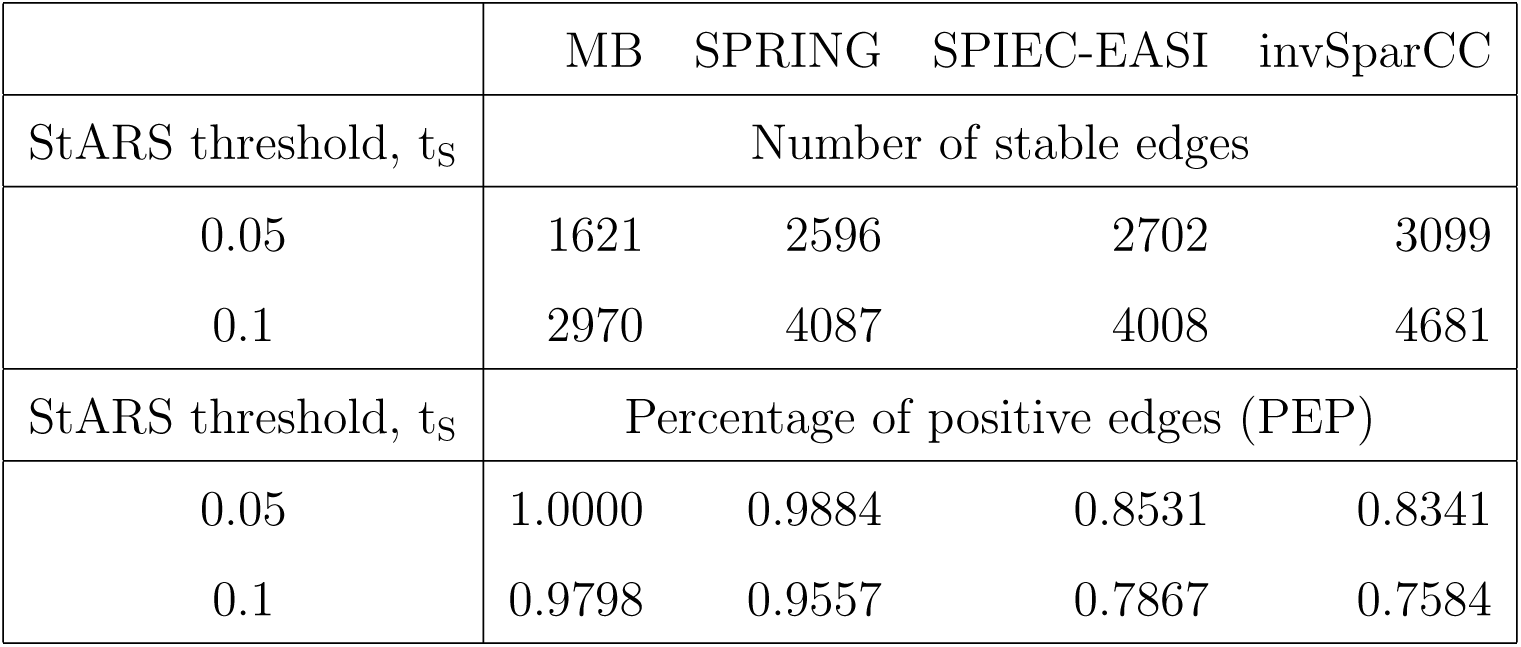
AGP data: Total number of partial correlation edges and percentage of positive partial correlation edges (PEP) (Faust et al., 2015) as estimated by MB, SPRING, SPIEC-EASI, and invSparCC for StARS stability thresholds t_S_ = 0.05 and 0.1.

|  | MB | SPRING | SPIEC-EASI | invSparCC |
| --- | --- | --- | --- | --- |
| StARS threshold, $t_s$ | Number of stable edges | | | |
| 0.05 | 1621 | 2596 | 2702 | 3099 |
| 0.1 | 2970 | 4087 | 4008 | 4681 |
| StARS threshold, $t_s$ | Percentage of positive edges (PEP) | | | |
| 0.05 | 1.0000 | 0.9884 | 0.8531 | 0.8341 |
| 0.1 | 0.9798 | 0.9557 | 0.7867 | 0.7584 |

To get a bird’s eye view of the topologies of the different association networks we visualize the four different networks at StARS threshold 0.05 in Figure 5 (A). The force-directed layout of all networks follows the optimal layout of the SPRING network. At the selected StARS threshold, all networks have one connected component. The overall network structure suggests a dense core with two peripheral network modules, similar to previous analysis (Müller et al., 2016). The networks of the compositionally-adjusted methods SPIEC-EASI and invSparCC connect the core and one of the modules by a large number of positive (shown in green) and negative (shown in red) associations. SPRING considerably sparsifies these connections, leaving only few positive and negative edges between the modules, and MB does not infer any negative associations. We assess the similarity among the estimated networks by analyzing their edge set overlap in Figure 5 (B). All methods share common core of 597 edges. As expected, SPIEC-EASI and invSparCC share the largest unique two-set overlap with 656. SPRING’s network takes an intermediate role between MB and the compositionally-adjusted methods. It shares 814 edges with SPIEC-EASI and invSparCC, and 112 edges exclusively with MB. Each method by itself also comprises a considerable set of exclusive edges, ranging from 437 for SPIEC-EASI to 694 for SPRING.

**Figure 5:**
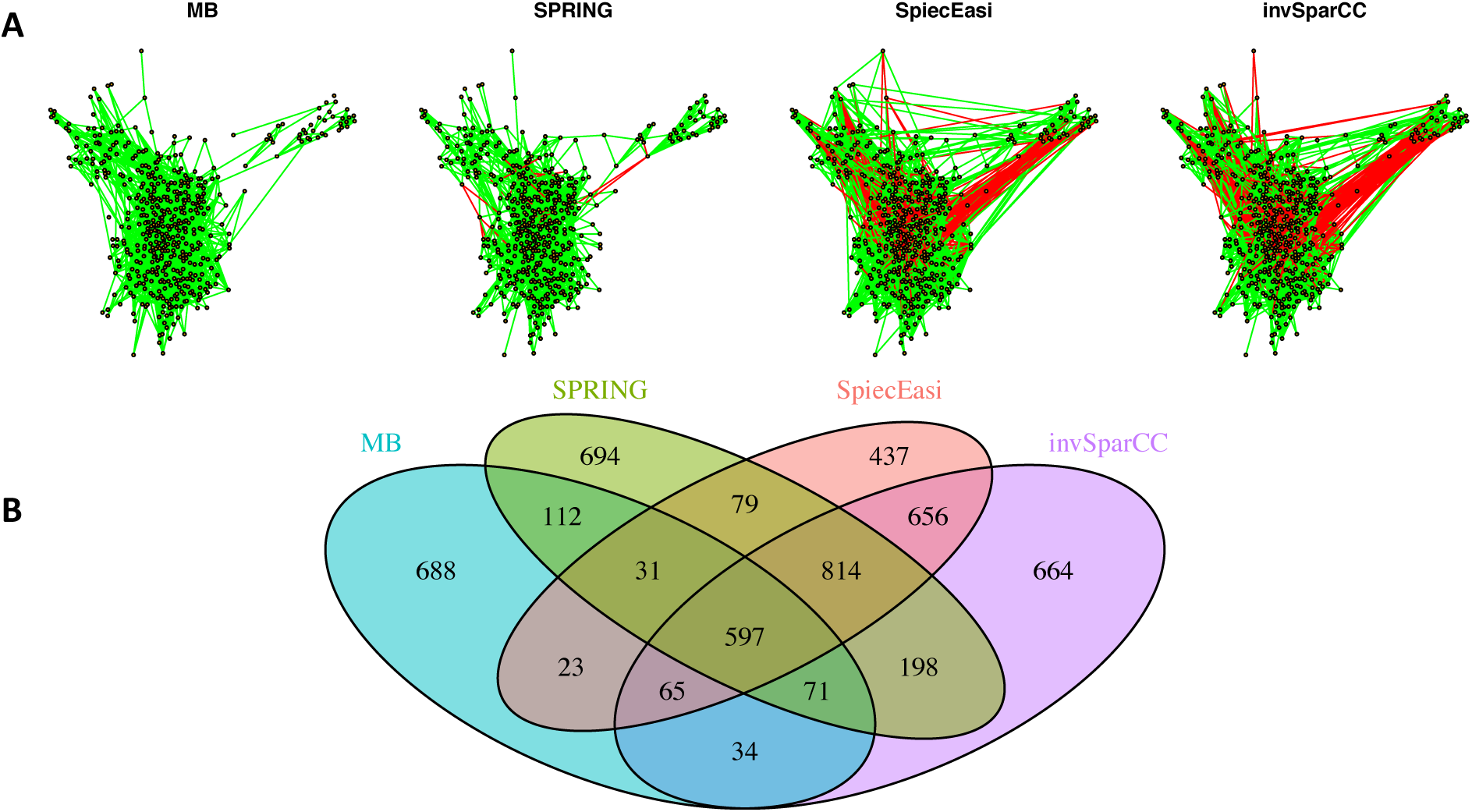
Analysis of AGP data. Panel (A): Force-directed layout (using igraph package in R) of the microbial association networks inferred by MB, SPRING, SPIEC-EASI, and invSparCC at StARS stability level 0.05. Green edges represent positive associations, red edges negative ones. Panel (B): The four set Venn diagram showing the overlap of edge sets of the different association networks.

### 4.2 Genus-genus associations from quantitative gut microbiome profiling data

We next analyze the quantitative gut microbiome data put forward in Vandeputte et al. (2017). We focus on estimating genus-genus associations both from the quantitative and the relative microbiome profiles, referred to as QMP and RMP, and analyze the consistency among the inferred networks. As the processed data used in Vandeputte et al. (2017) are not publicly available, we use the work flow outlined in Figure 1. We reprocessed the available amplicon sequencing data using the standard QIIME protocol with closed-reference OTU picking (Caporaso et al., 2010), adjusted for copy number variations of the 16S rRNA gene using PICRUSt (Langille et al., 2013), filtered the data as outlined in Section 3.2, and combined the samples with the measured total cell counts (Vandeputte et al., 2017). We pooled *n* = 106 healthy subjects from the two available cohorts and combined all OTUs on the genus level, resulting in *p* = 91 genera. To infer statistical genus-genus associations we use SPRING for the QMP data (without transformation), and SPIEC-EASI for the corresponding RMP data (using the standard clr transformation) with the same computational protocol as detailed in the previous section.

We first show the agreement of signed edges between the two association networks at StARS stability level 0.1 in Table 5. Overall, out of the 4095 possible genus-genus associations, SPRING infers a set of 237 stable edges with a PEP of 98%. SPIEC-EASI infers 220 edges with a PEP of 66%. From the quantitative data, SPRING is able to detect considerably more positive associations, 140 of whose are missed by SPIEC-EASI from the relative abundance data. SPRING also detects only four negative associations where three of those are missed by SPIEC-EASI despite having a considerable larger set of negative edges (74 overall). However, both methods do agree on a set of 93 edges, 92 positive and one negative edge. Importantly, we do not observe any sign flips among the different inferred edge sets. Missed positive or negative edges are simply absent in the other method.

**Table 5.**
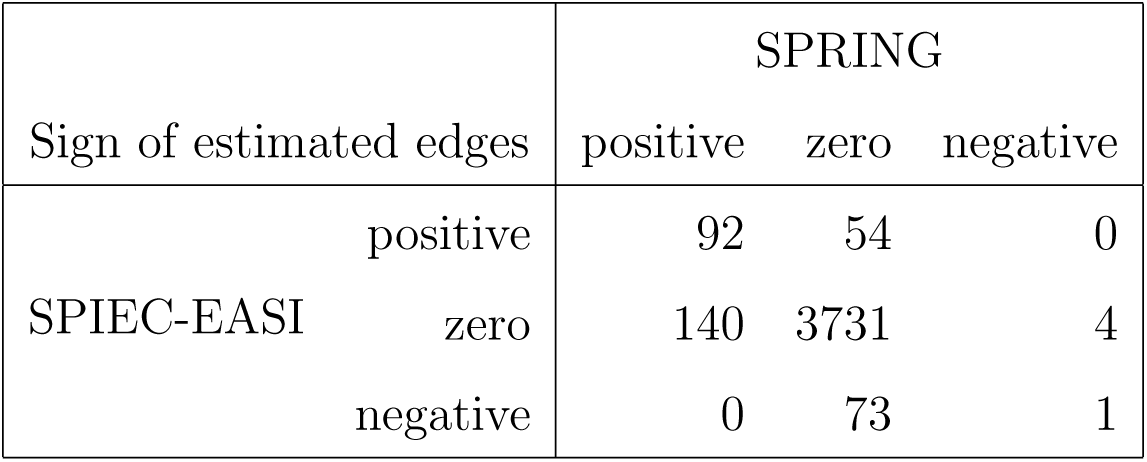
QMP data: Summary of agreement of signed genus-genus partial correlations, inferred by SPRING and SPIEC-EASI at StARS stability threshold t_S_ = 0.1.

We next focus on the induced genus-genus sub-network which only includes genera that have an assigned taxonomy and have at least one strong association *≥ |*0.2*|* in either the SPRING-inferred or SPIEC-EASI-inferred association network. The weighted adjacency of this sub-network includes 32 genera and is shown in Figure 6. Among the 14 genera with highest total abundance across all samples (Bacteroides to Odoribacter), we observe 50% agreement between the two estimated networks (six edges are the same across all networks, three edges are different in SPIEC-EASI, four are different in SPRING). Both networks include a strong negative association between Phascolarctobacterium and Dialister and exactly four positive associations of Bacteroides with Parabacteroides, Holdemania, Bilophila, and Odoribacter (First row and column in Figure 6). We also observe the absence of a negative association between Bacteroides and Prevotella genera in the quantitative data which is often reported in the literature and also present in the SPIEC-EASI network (see also Vandeputte et al. (2017) for a discussion).

**Figure 6:**
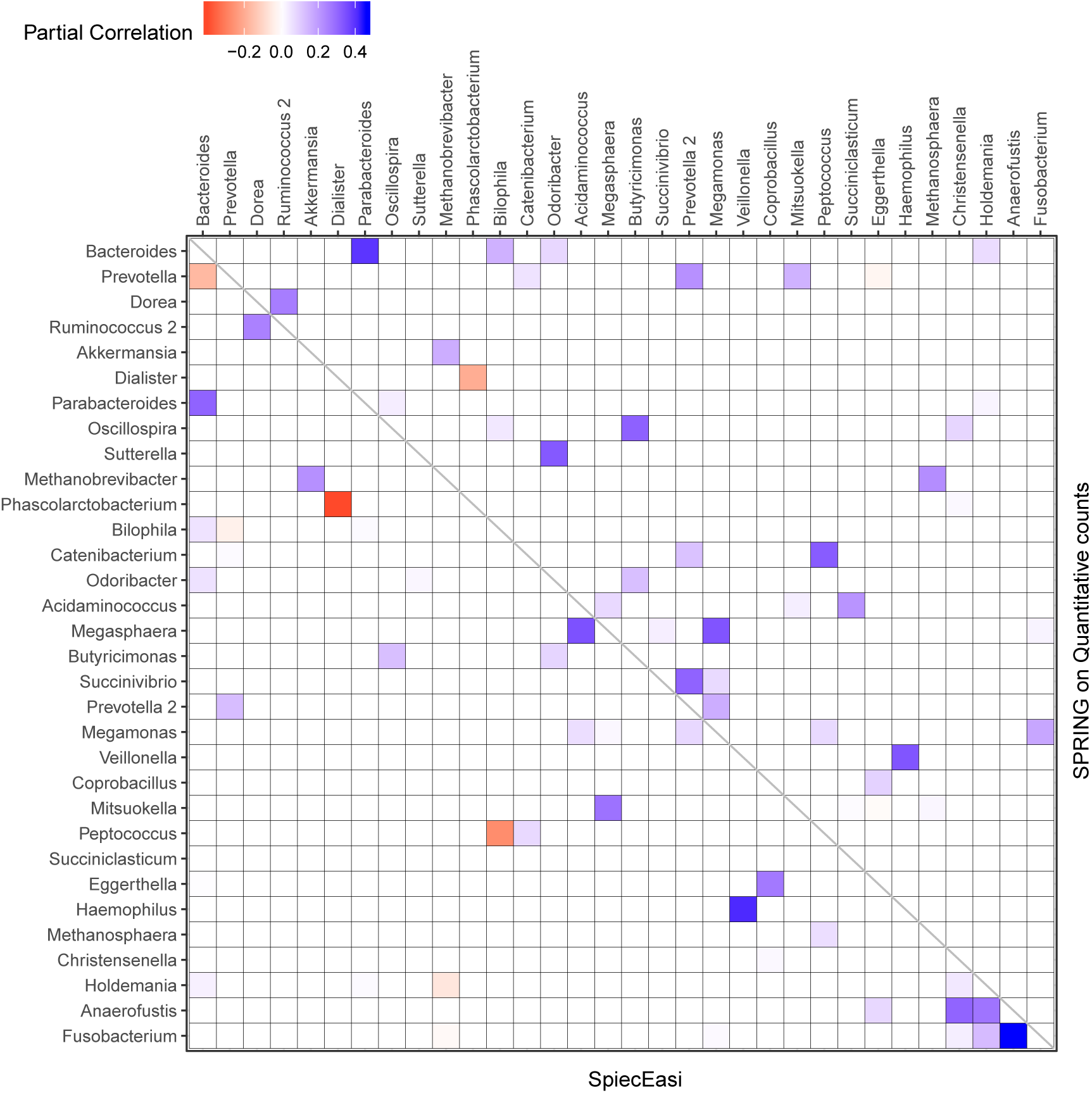
Genus-genus association network using relative (lower trianglur part) versus quantitative count data (upper triangular part). Only genera with at least one strong association ≥ |0.2| in either SPIEC-EASI or SPRING are shown. The genera are ordered by the total quantitative abundance over healthy subjects (*n* = 106).

## 5 Discussion

Advances in experimental microbiome profiling protocols have combined high-throughput environmental sequencing techniques with robust measurements of microbial cell counts (Gifford et al., 2011; Props et al., 2017; Satinsky et al., 2013; Stämmler et al., 2016; Tkacz et al., 2018; Vandeputte et al., 2017), providing, for the first time, a more quantitative picture of the underlying microbial ecosystems in their natural habitat. To facilitate a high-level summary of the complex interplay between the constituents of the ecosystem, an important first exploratory analysis step is the estimation of statistical association networks between the identified operational taxonomic units or gene sets (Faust and Raes, 2012; Fuhrman et al., 2015; Ruiz et al., 2017; Sunagawa et al., 2015). In order to learn such association networks from sparse quantitative microbiome data, we have introduced the **S**emi-**P**arametric **R**ank-based approach for **IN**ference in **G**raphical model (SPRING). SPRING combines neighborhood selection (Meinshausen and Bühlmann, 2006) to infer the conditional dependency graph with stability-based model selection (Liu et al., 2010; Müller et al., 2016) to identify a sparse set of partial correlation estimates. The resulting network of partial correlations represents direct (i.e., conditionally independent) microbe-microbe associations and provides a statistical community-level description of the underlying microbial ecosystem. As ground truth microbial association networks are largely elusive in the literature, we have based our numerical simulation benchmarks on a novel synthetic quantitative microbiome data generation mechanism which might be of independent interest to researchers who want to test novel statistical techniques on such data.

Our benchmark test cases revealed a number of interesting observations. First, we showed that naïve Pearson sample correlation estimation cannot be used to identify sparse partial correlations in quantitative microbiome data. Secondly, SPRING outperformed sparse graphical modeling techniques that were designed with compositional data in mind, namely SPIEC-EASI (Kurtz et al., 2015) and the invSparCC estimator introduced here, which uses neighborhood selection with SparCC correlation estimation (Friedman and Alm, 2012). SPRING compared favorably to the other methods both in terms of achievability, that is, in terms of minimum Hamming distance to the true underlying network achieved across the regularization path (see Fig. 3), and in combination with stability-based model selection in terms of Precision-Recall (see Fig. 4). We also quantified the robustness of SPRING to misspecification of the total by randomly distorting the counts of each sample up to a six-fold change which represents a realistic distortion scenario in gut microbiome samples (see e.g., in Vandeputte et al. (2017), Fig. 2) and is on the same order as typical fold changes of observed image-based total species counts in marine ecosystems (Fuhrman, 2018). Even under these distortions SPRING’s performance was on par or superior to SPIEC-EASI and invSparCC (which are scale-invariant by design). SPRING’s robustness to total count mis-specifications thus suggested to include an application of association inference from relative microbiome profiling data. In order to apply SPRING to relative abundance data we introduced a modified centered log-ratio (clr) transform that can seamlessly handle excess zeros without pseudo-count addition. Contrary to recent efforts in data-driven pseudo-count inference (see de la Cruz and Kreft (2018) and references therein) we computed the geometric mean of each sample from positive proportions only, normalized and log-transformed all non-zero proportions by using that geometric mean, and applied an identical shift operation to all non-zero variables in the dataset. This transformation is rank-preserving while leaving the original zero proportions unchanged, thus enabling the application of the SPRING methodology without further modification to relative abundance data.

We applied SPRING to two prominent gut microbial datasets, the relative abundance data collected in the American Gut Project (AGP) (McDonald et al., 2018) and the quantitative gut microbiome profiling (QMP) data from Vandeputte et al. (2017). As the processed data from Vandeputte et al. (2017) was not publicly available, a reprocessing of the amplicon sequencing reads was necessary.

From the AGP data, we inferred taxon-taxon association networks across *p* = 481 taxa from *n* = 6482 samples using neighborhood selection (MB), SPIEC-EASI, invSparCC, and SPRING. In line with previous findings (Faust et al., 2015), the percentage of positive edges in the networks is *>* 75%, with MB and SPRING having even higher percentages than SPIEC-EASI and invSparCC. At both StARS stability levels 0.05 and 0.1 reported here, SPRING and MB tended to infer slightly sparser association networks than SPIEC-EASI and invSparCC. At StARS stability level 0.05, we analyzed the overlap of edge sets among the different methods Fig. 5. All methods share a common core of 597 edges. In addition, SPRING, SPIEC-EASI, and invSparCC shared the largest common edge set of size 814 among all three-set overlaps. As expected, the two compositionally-adjusted methods SPIEC-EASI and invSparCC shared the largest common two-set overlap of 656 edges. In the absence of verified taxon-taxon associations, our analysis suggests that a practitioner screening for coherent statistical associations among taxa can apply SPRING, SPIEC-EASI, and invSparCC independently and select the set of strongest edges out of the edge set these three methods inferred. For the analysis here, this would result in 1470 edges, an average of three associations per taxon. This core network can then be further studied in terms of modularity, network stability, and node centrality measures, as shown, e.g., in Ruiz et al. (2017); Tipton et al. (2018).

For the QMP data, we used SPRING and SPIEC-EASI to estimate the genus-genus associations from the quantitative and the relative microbiome profiles, respectively. Our analysis revealed considerable differences to the published results in Vandeputte et al. (2017). The original study described dramatic differences between significant marginal genus-genus correlations from 66 healthy control samples in the QMP disease cohort when applying Spearman’s *ρ* correlation to the relative and quantitative microbiome profiling data (see, e.g., Fig. 3 in Vandeputte et al. (2017)). Our results here showed more coherence of the statistical associations inferred from relative and absolute abundance data. Overall, 92 positive, 1 negative, as well as 3731 zero associations were in common among both association networks, while both networks differed in 280 associations (Table 5). Our analysis on the genus sub-network that comprised all genera with at least one strong association *≥ |*0.2*|*, shown in Figure 6, verified a strong negative association between Phascolarctobacterium and Dialister inferred from both data types, as well as the absence of a negative association between Bacteroides and Prevotella genera in the quantitative data, both in agreement with published results. However, we recovered, for both data types, exactly four positive associations for Bacteroides, namely with Parabacteroides, Holdemania, Bilophila, and Odoribacter (First row and column in Figure 6). The latter two associations were previously reported only to be present in the quantitative data. Overall, more than 30% of the edges in the sub-network agreed which is in marked contrast to the results reported in Vandeputte et al. (2017). The higher network consistency reported here can be attributed to several factors. Firstly, our amplicon data processing framework may result in slight differences in terms of OTU picking and avoids a rarefaction step which was included previously. Secondly, we considered partial rather than marginal correlations among the genera to avoid any influence of indirect associations. Thirdly, we analyzed both data types within the same coherent statistical learning framework: sparse learning of partial correlations via neighborhood selection followed by stability-based model selection with the identical stability threshold (here 0.1). Finally, we considered a larger sample size of *n* = 106 representing healthy subjects from two different cohorts available in the QMP data as opposed to the *n* = 66 samples used in the original study. We conclude that differences in association networks from relative and absolute abundance data are not only attributable to the data themselves but also highly method-dependent.

In summary, we believe that, as quantitative microbiome profiling will become increasingly available, the semi-parametric rank-based estimators for correlation and partial correlation estimation discussed here provide an important tool for reliable statistical analysis of quantitative microbiome data. While we have focused here on targeted amplicon-based sequencing datasets, our methodology is broadly applicable to other biological high-throughput data with large excess of zero counts, including quantitative metagenomics (Satinsky et al., 2013), single-cell RNA-Seq data (see Risso et al. (2018) for a recent statistical analysis framework), and mass spectrometry proteomics data (Drew et al., 2017). Moreover, the concept of latent correlation employed in SPRING can naturally generalize to joint analysis of multiomics dataset when, on the same sample, several zero-inflated data types are measured in tandem. The approach in Yoon et al. (2018) already exploits this idea for RNA-seq and micro-RNA data in the context of canonical correlation analysis. Extending SPRING in a similar way to joint graphical modeling of mixed data types is a promising next step toward a consistent and coherent statistical analysis framework for sparse high-throughput biological datasets.

## Author Contributions

GY, IG and CLM developed the methodology; GY lead the numerical analysis; IG assisted GY in simulation studies; CLM assisted GY in data analyses; GY, IG and CLM prepared the manuscript.

## Funding

GY was supported by the National Cancer Institute grant T32-CA090301. IG was supported by the National Science Foundation grant DMS-1712943. CLM work was supported by the Simons Foundation.

### Acknowledgments

We are grateful to Dr. Zachary D. Kurtz, Lodo Therapeutics, for providing the processed American Gut data and the QMP data.

## Supplemental Data

### Data Availability Statement

Data from the American Gut Project are available at https://github.com/biocore/American-Gut. The raw amplicon sequencing data for the QMP study are deposited in European Nucleotide Archive with accession codes PRJEB21504 and ERP023761.

